# Human intestinal organoids with an autologous tissue-resident immune compartment

**DOI:** 10.1101/2023.10.04.560810

**Authors:** Timothy Recaldin, Bruno Gjeta, Linda Steinacher, Marius F. Harter, Lukas Adam, Mikhail Nikolaev, Rok Krese, Umut Kilik, Doris Popovic, Marina Almató-Bellavista, Kristina Kromer, Michael Bscheider, Lauriane Cabon, J. Gray Camp, Nikolche Gjorevski

**Affiliations:** Institute of Human Biology (IHB), Roche Pharma Research and Early Development; Roche Innovation Center Basel, Roche Pharma Research and Early Development; Gustave Roussy Cancer Campus, University Paris-Saclay, Paris, France; University of Basel, Basel, Switzerland; Hannover Medical School, Institute of Immunology, Hannover, Germany

**Author notes:** Correspondence: J. Gray Camp & Nikolche Gjorevski.

## Abstract

The intimate relationship between the epithelium and the immune system is crucial for maintaining tissue homeostasis, with perturbations in epithelial-immune interactions linked to autoimmune disease and cancer. Whereas stem cell-derived organoids are powerful models of tissue-specific epithelial function, these structures lack tissue-resident immune cells that are essential for capturing organ-level processes. We describe human intestinal immuno-organoids (IIOs), formed through self-organization of epithelial organoids and autologous tissue-resident lymphocytes (TRMs), a portion of which integrate within the IIO epithelium and survey the barrier. IIO formation was driven by TRM migration and interaction with epithelial cells, as orchestrated by TRM-enriched transcriptomic programs governing cell motility and epithelial inspection. We combined IIOs and single-cell transcriptomics to investigate intestinal inflammation triggered by cancer-targeting biologics in patients, and found that the system recapitulates clinical outcomes and the underlying cellular mechanisms. Inflammation was associated with the emergence of an activated population of CD8+ T cells, which progressively acquired intraepithelial and cytotoxic features. The appearance of this effector population was preceded and likely mediated by a Th1-like CD4+ population, which initially displayed a cytokine-producing character and subsequently became cytotoxic itself. A system amenable to direct perturbation and interrogation, IIOs allowed us to identify the Rho pathway as a novel target for mitigating immunotherapy-associated intestinal inflammation. Given that they recapitulate both the phenotypic outcomes and the underlying inter-lineage immune interactions, IIOs can be used to broadly study tissue-resident immune responses in the context of tumorigenesis, infectious and autoimmune diseases.

## Main text

Organoids originating from adult stem cells (ASCs) model important aspects of human physiology, and have found applications in research related to genetic disorders, infectious disease, cancer, regenerative medicine and drug discovery (1–6). However, ASC-derived organoids are epithelial-only structures, whereas native organs comprise multiple other compartments, including highly specialized immune cells that interact with the epithelium and play essential roles in homeostasis and disease (7–11). For example, the intestinal mucosal immune system, which represents the largest pool of immune cells in the human body (7), ensures homeostasis by perpetually policing the interface between the intestinal barrier and luminal contents. Disruption of intestinal immune function can lead to various pathologies, including persistent infections, autoimmune disorders and even malignant disease (12–17). Whereas intestinal organoids can accurately model many aspects of the differentiation and function of epithelial cell types (18–21), they fall short in capturing essential aspects of intestinal homeostasis and disease, owing to the absence of a functional and tissue-specific immune compartment (22, 23).

In order to address these shortcomings, adult human intestinal epithelium has been cocultured with blood-derived innate immune cells (24, 25). However, the incorporation of a mucosal lymphocyte compartment has proven elusive. A recent study described the generation of gut-associated lymphoid tissue in iPSC-derived human intestinal organoids upon transplantation in humanized mice (26). Although an important advance, the in vivo formation of immune tissue removes some of the advantages of organoids as controllable in vitro systems. Further, owing to its iPSC origin, this system approximates a fetal intestine, rather than an adult one.

Here, we created a tractable intestinal immuno-organoid (IIO) model containing a tissue-resident and autologous immune compartment starting from readily available adult human tissue samples (Fig. 1a). We benchmarked cell states in IIOs through comparison to reference atlases using single-cell transcriptomes, and used IIOs to recapitulate and investigate drug-induced intestinal inflammation. In our efforts to introduce a functional and relevant lymphocyte compartment into intestinal organoids, we focus on tissue-resident memory T cells (TRMs). TRMs are antigen-experienced T-cell populations, which take permanent residence in the intestinal mucosa, providing front-line defense against invading pathogens (10, 11). Given the absence of recirculation, they are an appropriate lymphocyte type to stably incorporate into organoid models. Furthermore, their prior antigen exposure and memory character ensures functionality in the absence of antigen-presenting cells, lymphoid structures and the remaining immune cell recirculation machinery. TRMs are difficult to incorporate into in vitro systems, owing to their poor viability upon enzymatic removal from the tissue (27, 28). Therefore, we adapted an enzyme-free scaffold-based crawl-out protocol to isolate large numbers of healthy intestinal immune cells (29) (Fig. 1a). We found that, even in the complete absence of cytokine or T-cell receptor (TCR) stimulation, our approach liberated significantly more cells than enzymatic digestion-based protocols, while retaining similar proportions of immune cell types (Extended Data Fig. 1a-b). We deemed this lack of cytokine exposure crucial for retaining the tissue-like physiological properties of the intestine-derived lymphocytes. Indeed, flow cytometry analysis showed that the isolated cells expressed TRM markers pertinent to the intestine, including CD161 (IL-17A production (30)) and CD117 (notch signaling (31)) (Extended Data Fig. 1c), as well as surface molecules associated with tissue retention (CD69 (32)), extracellular matrix association (CD49a (33)) and epithelial cell integration (CD103 (34)), all of which were undetectable on blood-derived lymphocytes (Fig. 1b-c).

**Figure 1.**
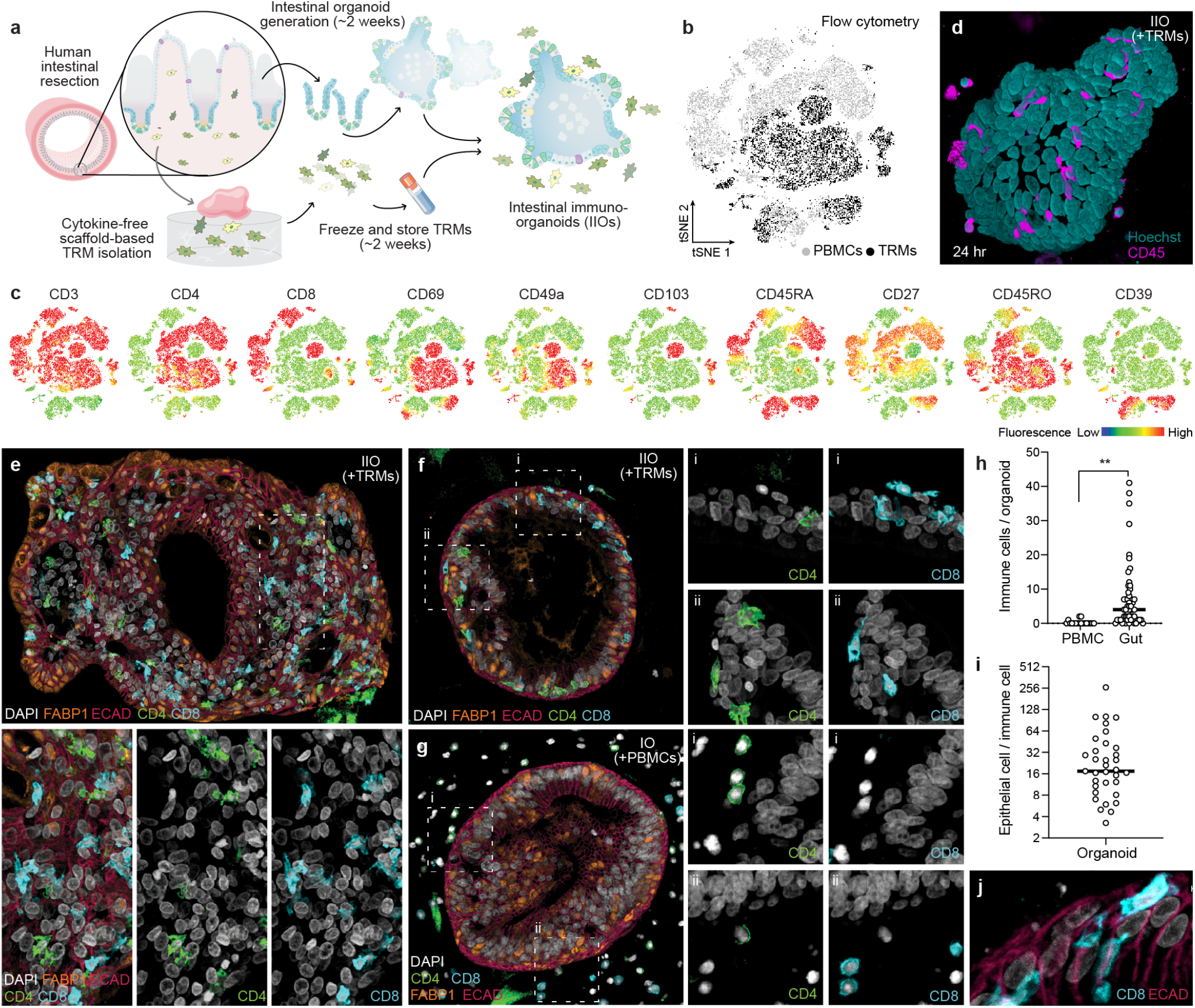
Intestine-derived tissue-resident lymphocytes (TRMs) integrate homeostatically into autologous organoids to form intestinal immuno-organoids (IIOs). a, Schematic overview for establishing autologous IIOs. b-c, Flow cytometry-based tSNE analysis of gut (TRM) and circulating (PBMC) T cell subgroups based on surface marker expression. tSNE plot colored by original source of the T cells (b, light grey for PBMC and dark grey for TRM) and expression of 10 individual markers of naivety, memory or tissue residency (c). d, Fluorescent still from live imaging 24 hours after IIO coculture with autologous TRMs (nuclei = teal, T cells = pink). e-g, Fluorescent IHC staining of IIO cultures 24 hours after coculture with autologous TRMs (e-f) or PBMCs (g). h, Detected immune cell count per organoid, each data point represents 1 individual organoid. **P<0.01, unpaired T-test. i, Ratio of epithelial cells to immune cells within each organoid, following organoid supplementation with autologous TRM cells. j, Elongated flossing T cell inserting between basal-lateral epithelial cell junctions.

We generated organoids and TRMs from human intestinal specimens, additionally collecting matched peripheral blood mononuclear cells (PBMCs) from the same donor. Once established, organoids were combined with TRMs or PBMCs within three-dimensional (3D) extracellular matrix (ECM) at physiologically-relevant cellular concentrations (35), and in the absence of external stimulation. Confocal microscopy and 3D reconstruction revealed that, after 24h coculture, TRMs were viable and closely associated with the organoids (Fig. 1d, Extended Data Video 1). To examine the organoid-TRM interactions in greater detail, we generated histological sections of the models and visualized the epithelial and immune cells (Fig. 1e-g). We found that, whereas PBMCs occupied the ECM space without apparent interactions with the epithelial cells (Fig. 1g), a sub-population of TRMs infiltrated the organoids and integrated within the epithelial barrier, strikingly resembling the behavior of intestinal intraepithelial lymphocytes (IELs) (36) (Fig. 1e-f, h). We estimated a median integration ratio of 16 epithelial cells per immune cell (Fig. 1i) - highly similar to observations in the intestinal tract of healthy humans (37). Intraepithelial lymphocytes displayed an elongated morphology of 60 pm in length (Fig. 1j, Extended Fig. 1d), around 10 times the length of a naive blood-derived T cell (38), and reminiscent of the “flossing” behavior described for transgenic murine lELs imaged in vivo (36). With low-level cytokine support, immune-organoid cultures could be maintained across at least three passages, albeit with a diminishing ratio of immune cells to epithelial cells (Extended Data Fig. 1e). This model provides the first example of self-organization between human immune cells and epithelial organoids to form an organoid system with a tissue-resident immune compartment. We termed these structures intestinal immuno-organoids (IlOs).

Elegant in vivo studies have dissected the dynamics of murine TCRyb lELs and their interaction with the intestinal epithelium (36). Owing to their poor survival in vitro, similar studies of human IEL behavior have been challenging (27, 28). As a result, the mechanisms driving human-specific IEL integration are poorly elucidated. To understand how TRMs and lELs are capable of interaction and integration with intestinal epithelial cells in vitro, and how they differ from PBMCs in that regard, we used single-cell RNA sequencing (scRNA-seq) to analyze donor-matched tissue-resident and blood-derived immune cells alone, or cocultured with organoids (Fig. 2a). After dissociating the cultures, we prepared scRNA-seq libraries of TRMs and PBMCs, with a focus on CD3+ T cells, given the TRM predominance within the tissue-derived population. Heterogeneity analysis and visualization using UMAP embedding demonstrated the presence of 3 distinct populations, representing blood-derived naive, blood-derived memory and gut-derived tissue-resident memory T cells (Fig. 2b, Extended Data Fig. 2, Extended Data Table 1). Further interrogation of the T cell populations based on previously published markers (39–42) revealed that TRMs, unlike their matched blood-derived counterparts, were transcriptomically defined by: (i) the absence of receptors necessary for lymph node homing (SEL, CCR7), indicative of their non-circulating tissue-residence status, (ii) intrinsically high expression of intestinal integration factors (ITGA1, CCR9, JAML), and (iii) a complete absence of cytotoxic granules (GZMA, GZMB, GNLY, NKG7) despite their function as highly differentiated effector T cells (Fig. 2c, Extended Data Table 2). We observed that populations c1, c3 and c8 were uniquely enriched in gut-derived TRMs (Fig. 2d). To identify the main functional differences between TRMs and PBMCs, we considered the top differentially regulated TRM genes, and observed a dominance of genes governing cell adhesion, motility and cytoskeletal rearrangements (Fig. 2e). Gene ontology analysis suggested enrichment in transcriptomic programs related to immune cell chemotaxis and migration within TRMs compared with PBMCs (Fig. 2f), which may explain the different propensities for movement toward and integration within the epithelium between these two populations. Indeed, live imaging experiments in which donor-matched TRMs and PBMCs were tracked over time (Fig. 2g; Extended Data Video 2) showed a striking difference in morphologies and migratory behaviors between the two populations. Whereas PBMCs were static and morphologically round, TRMs exhibited asymmetric, elongated shapes with front-rear polarity, and migrated dynamically within both the epithelial layers and ECM, with speeds of up to 80 pm/h (Fig. 2h). TRM migration was not directional, suggesting that TRMs in this system integrate within organoids through random movement and encounters of epithelial cells, rather than through chemotaxis (Extended Data Video 3). Indeed, TRMs appeared to be as likely to move towards and within organoids as they were to move out and away. We note that intestinal organoids are a sterile system and the introduction of luminal microbes may lead to altered modes of migration and interaction with the epithelium, as described in mice (36).

**Figure 2.**
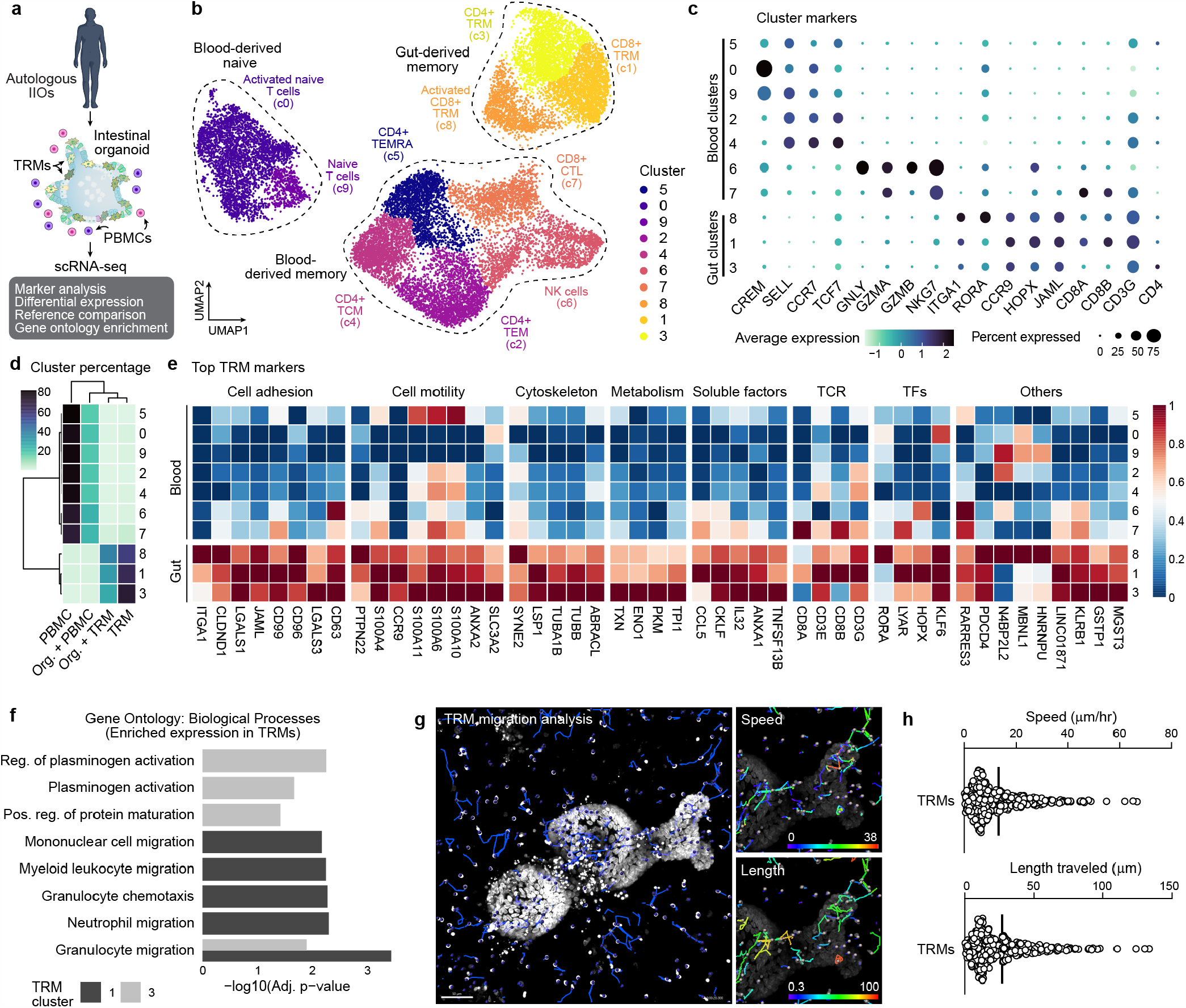
Tissue-resident transcriptomic signatures and migratory behavior underlie TRM epithelial insertion and IIO formation. a, Overview of single-cell transcriptome analysis of IIOs containing patient-matched TRMs and PBMCs. b, Integrated UMAP embedding of transcriptome data shows 10 distinct cell clusters (colors, numbers) labeled based on analysis of marker genes. c, Dotplot summarizing marker gene expression across the different clusters. d, Heatmap representing proportional sample distribution in each cluster. e, Heatmap representing TRM-specific genes grouped by functional category. TFs transcription factors, TCR T-cell receptor. f, Barplot showing significantly enriched gene ontology (GO) biological processes of genes marking TRM clusters 1 (CD8+) and 3 (CD4+). g, Images showing migration analysis from IIO time-lapse imaging. Tracks are shown in blue (left) and colored by speed (top left) and length (bottom right). Nuclei are labeled with Hoechst (white). h, Distribution of TRM speed (top) and length traveled (bottom) from IIO time-lapse imaging.

Next, we tested whether differential transcriptomes and migration behaviors between TRMs and PBMCs translated into differences in effector function within IlOs. In particular, we investigated whether IIOs can recapitulate clinical toxicities associated with cancer immunotherapy that manifest as severe intestinal inflammation and were not predicted by conventional preclinical models (43, 44). We focused on solitomab - a bispecific T-cell engager intended to crosslink activated T cells to solid tumors via EpCAM - which induced rapid and aggressive unintentional intestinal inflammation in patients, leading to program termination (44). Given the rapid onset of the side effects, IELs localized within the basolateral epithelial junctions have been hypothesized to elicit the damage (44). To assess whether IEL-containing IIOs could have predicted targeting of the healthy epithelium, we treated IIOs with an EpCAM-targeting T-cell bispecific (TCB) molecule at concentrations relevant to those detected in the serum of patients of the solitomab trial (44). Unlike organoids cultured with PBMCs, IIOs were aggressively targeted in a TCB-dose-dependent manner at concentrations as low as 40 pg/mL, as demonstrated by the detection of caspase 3/7 (Fig. 3a-c). A time-course of epithelial cell targeting showed induction of caspase by TRMs as early as 8 h after treatment (Extended Data Fig. 3a), mirroring clinical observations that were not predicted with classical in vivo models. We assessed T cell behavior at early (5h), mid (24h) and late (48h) timepoints by digesting and staining IIOs for surface and intracellular markers of T-cell activation and cytotoxicity (Extended Data Fig. 3b). Identification of effector populations, based on the expression of TNFa, IFNy and granzyme B, revealed that over 90% of responding cells were intestinal TRMs (Extended Data Fig. 3c). This demonstrated that TRMs were approximately 10 times as likely to induce a detectable response to TCB treatment than their blood T-cell counterparts, emphasizing the necessity to include appropriate cell types within in vitro tissue-specific models. The mechanisms of severe rapid-onset toxicities caused by T-cell targeted therapies are unclear, as patients who experience them are not biopsied in the acute phase. IIOs, which we show can recapitulate clinical outcomes, provide the opportunity for in-depth analysis of the underlying cellular and molecular events. We used scRNA-seq to interrogate the transcriptomic dynamics underlying TCB-dependent TRM activity at the onset (4h) and peak (48h) of epithelial cell targeting. Lymphocyte populations within the integrated dataset were annotated using differential gene expression together with previously published signatures and surface markers (39–42, 45), revealing diverse T cell, macrophage and B cell populations (Fig. 3d-e, Extended Data Table 1 and 2). TCB treatment induced a time-dependent shift in proportions of both the T cell and B cell states relative to the non-targeting control (Fig. 3f). This transition was driven by the emergence of several effector populations at both time points. Particularly prominent at 4h was a CD4+ T helper 1-like (Th1) population (c9) (Fig. 3f), characterized by a rapid induction of TNF and IFNG signaling that became down-modulated over time (Fig. 3g-h). At 48h, we observed the emergence of an activated CD8+ IEL population (c5) (Fig. 3d, f), expressing genes related to cytotoxicity (such as GZMB), TCR signaling and T cell migration (Fig. 3g-h). Concurrently, a population of cycling (MKI67-expressing) CD4+ T cells (c12) and an activated population of B cells (c11) appeared, whereas the Treg population (c1) diminished (Fig. 3d, f, g). Importantly, key gene expression changes detected by scRNA-seq were mirrored by changes at the protein level, obtained by flow cytometry (Fig. 3i). In particular, we observed early induction of TNFa in CD4+ cells at 4h (corresponding to c9), whereas the cytotoxic granzyme B was upregulated at the 48h time point, consistent with the appearance of c5. Likewise, Ki67 was strongly upregulated within a subset of CD4+ cells at 48h, co-occurring with the appearance of c12 within the scRNA-seq analysis. IIO cell heterogeneity dynamics bear striking similarities to those observed within primary samples from patients experiencing spontaneous and immune checkpoint inhibitor (ICQ-induced intestinal inflammation. For example, the emergence of a cytotoxic CD8+ T-cell population was one of the main features of patients experiencing ICI-induced colitis (40, 46). Likewise, Treg depletion and the appearance of an activated IFNy-responsive B-cell population were reported in clinical samples from colitis patients (47, 48). These similarities suggest that IIOs may be used to recapitulate and study intestinal inflammation in a tractable in vitro setting

**Figure 3.**
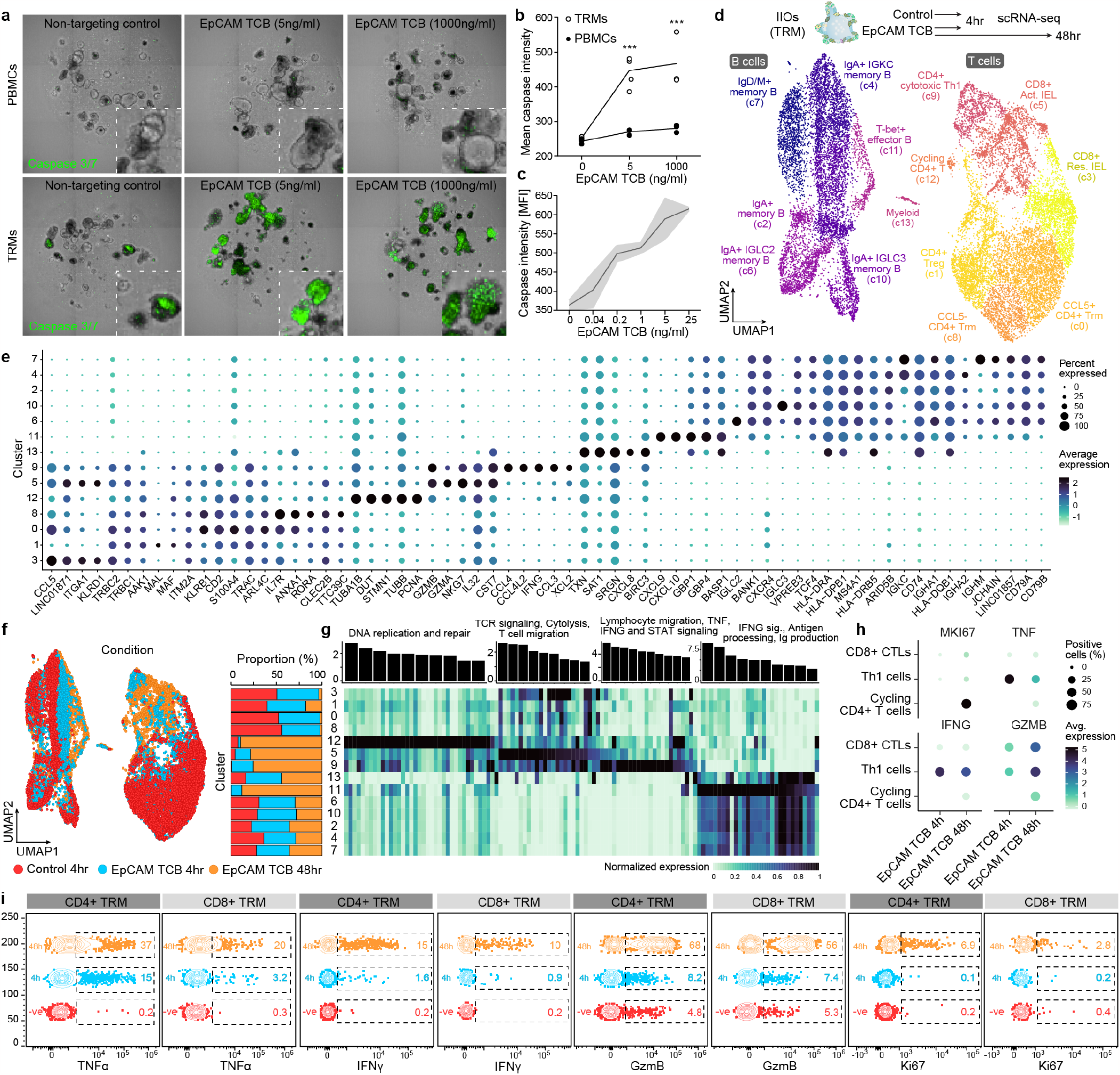
IIOs recapitulate clinically manifested intestinal inflammation associated with T cell bispecific antibodies (TCBs). a, Representative images examining induction of green caspase 3/7 signal within IIO cocultures 24h after supplementation with EpCAM TCB. b, Quantification of the caspase 3/7 signal from (a). ***P<0.001, 2way ANOVA with Sidak’s multiple comparisons test. c, Quantification of the caspase 3/7 signal in IIO cocultures treated for 72h across a range of EpCAM TCB, n = 3. d, Single cell transcriptomic profiles of gut-derived immune cells from IIO model were integrated and grouped into 14 distinct cell states as represented by the colors in the UMAP embedding. e, Dotplot summarizing the expression patterns of representative genes across the clusters identified in (d). f, Integrated UMAP embedding (left) and proportional distribution (right) of gut-derived immune cells from IIO model colored by treatment and profiling time. g, Barplot showing significantly enriched GO biological processes for activated cell states (top) and heatmap showing average expression profiles of corresponding associated genes (bottom). h, Dotplot summarizing the expression pattern of representative genes involved in proliferation, signaling and cytotoxicity in the activated T cell populations as captured by scRNA-seq snapshots at different timepoints and treatment conditions. i, Flow cytometry plots visualizing expression of TNFa, IFNy, GzmB and Ki67 across the different timepoints within CD4+ and CD8+ TRM cells isolated from IIO cultures.

Next, we set out to chart the dynamics of clinically relevant populations that appear in IIOs, focusing first on the progression of CD8+ T-cell activation. Using Diffusion Maps (49) we computed a pseudo-temporal ordering of CD8+ populations (c3 and c5) (Fig. 4a, Extended Data Fig. 4a) and observed a striking correlation with experimental time (Fig. 4b). This reconstructed activation trajectory allowed for characterization of the transcriptional dynamics underlying CD8+ TRM lymphocyte activation (Extended Data Fig. 4c). In particular, we observed strong induction of glycolytic regulators ENO1 and HIF1A, with concurrent suppression of TCF7 and ZBTB32 (50–53) to likely facilitate the appropriate metabolic profile for full CD8 TRM cell activity. Simultaneously, CCL5, important for immune cell recruitment and early inflammatory responses (54), and IL7R, which captures proinflammatory signals to mediate cytotoxic activation (55), correlated with induction of cytotoxicity effector molecules GZMA, GZMB and NKG7. Sequencing of inflamed colonic biopsies of patients suffering from drug-induced colitis demonstrated the presence of both cytotoxic (CTL) and IEL CD8+ T cell populations (40). By cross referencing TCB-treated IlOs to this dataset, we found that IIO CD8+ T cells acquired gene signatures related to a cytotoxic and IEL state, mirroring those observed in clinical samples (Extended Data Fig. 4d-e). The concurrent increase in lymphocyte-epithelium association and cytotoxicity may underlie the severe clinical adverse events triggered by these molecules.

**Figure 4.**
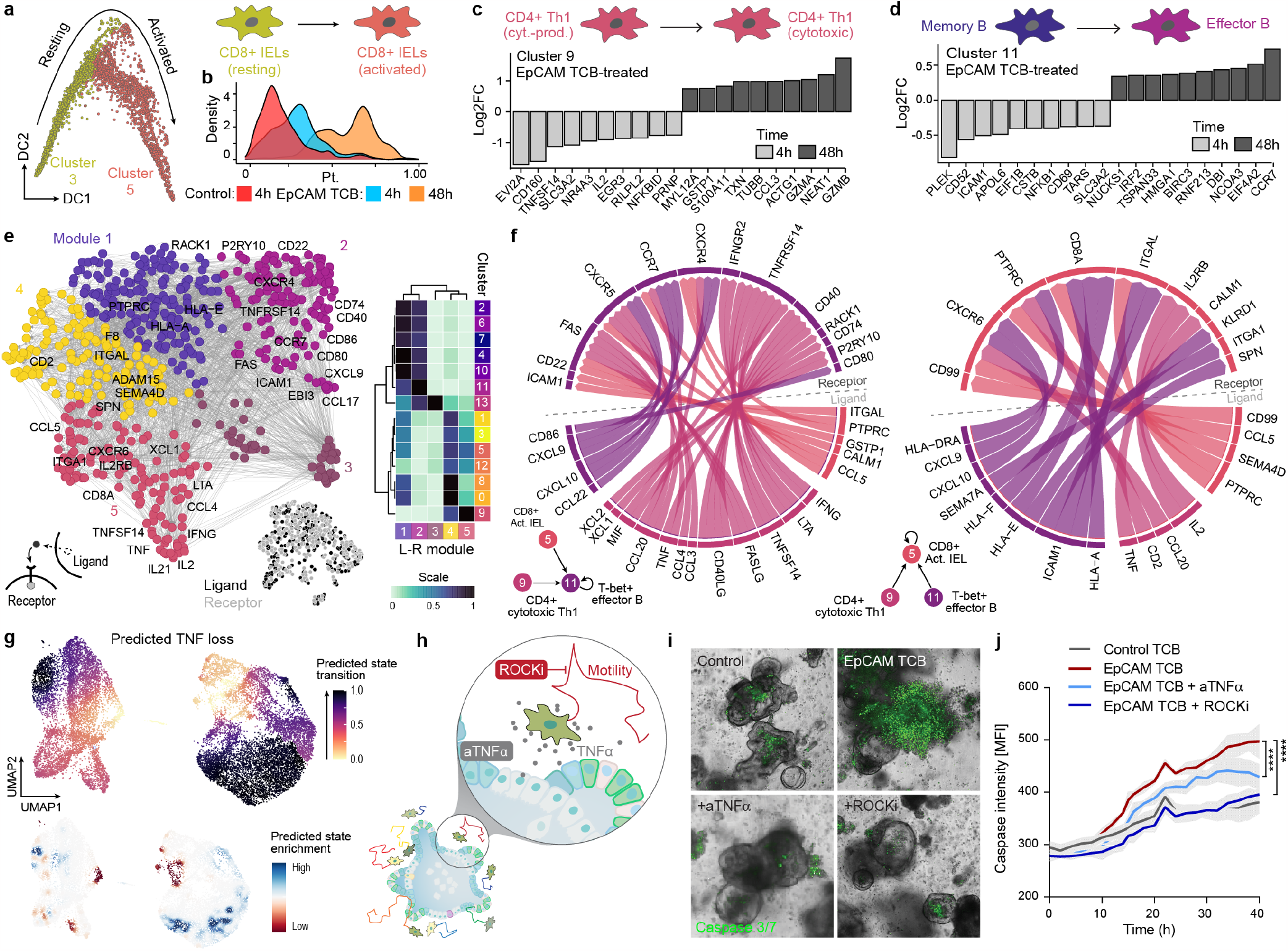
Transcriptomic analyses elucidate the immune dynamics underlying TCB-mediated inflammation and help identify mitigation strategies. a, CD8+ T cell activation in IIO model was analyzed with diffusion maps. Plot represents CD8+ T cells on the first two diffusion components colored by cell state. b, Density plot showing distribution of CD8+ T cells along the reconstructed pseudotime (x-axis) grouped by treatment condition and time point. c-d, Barplot showing differentially expressed genes for CD4+ cytotoxic Th1 population (c9) (c) and T-bet+ effector B cells (c11) (d) at 4h (light grey) and 48h (dark grey) after EpCAM TCB treatment. e, Ligand-receptor pairing analysis of IIO immune cell populations. Ligands and receptors are colored based on co-expression module, with some representative genes labeled. Heatmap (right) shows the average expression of each gene within a module across each cell cluster. f, Circle plots describe signaling interactions received by the T-bet+ effector B cells (c11) (left) and the CD8+ Act. IELs (c5) (right). g, In silico perturbation analysis simulating the loss (KO) of TNF in IIO model treated with EpCAM TCB. Plots show predicted perturbation-induced state transition (top) or enrichment (bottom). Note that the activated immune cell states are predicted to fall back to homeostatic conditions. h, Schematic of experiments to inhibit TNFa signaling and cell migration. i, Representative images examining induction of green caspase 3/7 signal within IIO cocultures 40h after supplementation with either a non-targeting control TCB or 5 ng/ml EpCAM TCB with or without 1 mM Y27362 (ROCKi) or a TNFa blocking antibody. j, Bihourly quantification of caspase 3/7 signal in IIO cultures in treatment conditions, n = 3. ****p < Q.QQQ1, T-test of the AUC compared to the EpCAM TCB condition.

Aside from CD8+ CTLs, other populations displayed clinically relevant dynamics of early-vs. late-transcriptional hallmarks. Consistent with clinical reports (56), a Th1 population (c9) shifted from a cytokine- (TNF, IFNG and IL2)-producing state to a cytotoxic, GZM-producing state (Fig. 4c, Extended Data Fig. 4f). Likewise, the early IFN-responsive B cell population (c11) showed a transcriptional landscape distinct from that of the late, activated state (Fig 4d, Extended Data Fig. 4g). We performed a receptor-ligand pairing analysis and inspected how activated phenotypes may emerge via intercellular signaling (Fig. 4e). Our model implicates Th1 cells (c9) as a major organizational hub, instructing B cells (c11) and CD8+ T cells (c5) through the secretion of numerous signaling factors (Fig. 4f, Extended Data Fig. 4h). In particular, chemokines (CCL3, CCL4, CCL5 and CCL20) and pro-inflammatory molecules (IFNG, CD40LG and TNFSF14) expressed by T cell clusters 5 and 9 likely encouraged close association with, and then full activation of, B cells via reciprocal receptors (Fig 4f, Extended Data Fig. 4i). Meanwhile, CD4 T cell (c9) expression of TNF corresponded to increased CALM1 expression in activated CD8+ IELs (c5), potentially augmenting TCR-induced calcium signaling and full T-cell maturation (57) (Fig. 4f, Extended Data Fig. 4j). Ligand-to-target signaling models and network propagation analysis (58) reaffirmed these observations, suggesting that Th1-produced IFNG may mediate the up-regulation of activation-related genes within B cells, while TNF and IL2 act in tandem to orchestrate cytotoxic CD8+ cell maturation (Extended Data Fig. 4h-i).

A key advantage of human model systems is their amenability to direct experimental manipulation as a means to define the roles of putative regulators. Given the function of TNFa as an instigator of inflammation, the effectiveness of TNFa-blocking antibodies in the treatment of autoimmune disease (59), and its prominent early induction in our model (Fig. 3h-i), we investigated the role of this cytokine in promoting differentiation and activation profiles. In silico perturbation analysis simulating the complete removal of TNF from IIOs predicted that TNF depletion would prevent the emergence of activated immune cell populations and ultimately reduce the cytotoxic response (Fig. 4g). Antibody-neutralisation of TNFa in IIOs confirmed these predictions, significantly reducing epithelial cell apoptosis following TCB treatment (Fig. 4h-j). Having confirmed the impact of neutralizing a clinically-validated pathway, we used the transcriptomic data defining TRM identity (Fig. 2) to suggest novel TRM-specific factors that could be manipulated to quell unwanted TCB-mediated inflammation. Given the TRM cytoskeletal transcriptomic signature and their rapid locomotion within the ECM, we hypothesized that T-cell motility and cytoskeletal rearrangements may be in part responsible for the outcome. To test this hypothesis, we used the ROCK1/2 inhibitor (ROCKi) Y27632 to abrogate cell motility (60) within TCB-treated IIOs. Strikingly, we found that ROCK inhibition reduced epithelial cell apoptosis even more efficiently than TNFa blockade (Fig. 4h-j), simultaneously suppressing the induction of T-cell activation markers such as CD25, as well as cytolytic molecules such as perforin and granzyme B (Extended Data Fig. 5). In addition to TNFa and Rho-mediated signaling, the transcriptomic and ligand-receptor analyses shown above provide a wide range of putative targets for managing inflammation, which are worth investigating in future studies.

We thus describe human intestinal immuno-organoids (IIOs), comprising human sample-derived intestinal epithelium and autologous tissue-resident lymphocytes (TRMs), a sub-population of which are directly integrated within the IIO epithelial barrier, reflecting IEL inclusion within the native intestine. IIO formation was driven by extensive TRM migration and interaction with epithelial cells, as orchestrated by TRM-enriched transcriptomic programs governing cell motility and adhesion. Crucially, IIOs formed upon coculture with TRMs, but not blood-derived lymphocytes, which lacked the migratory properties of the former and failed to interact with organoids.

The inclusion of a tissue-appropriate immune compartment extends the utility of organoids far beyond epithelium-centered questions and applications. We use IIOs to recapitulate intestinal inflammation caused by cancer-targeted bispecific antibodies (TCBs) in phase I clinical trials. IIOs treated with TCBs at clinically relevant concentrations undergo rapid apoptosis, consistent with early-onset diarrhea and epithelial lesions in patients (44). Importantly, whereas co-cultures with PBMCs have been shown to capture similar outcomes, effects become apparent only at concentrations 1000-fold higher than clinical doses and, even then, with a delayed onset [MFH, TR, MB, NG, manuscript in preparation]. Dissecting the transcriptomic changes induced by TCB treatment, we uncover that the adverse outcomes are underpinned by elaborate and dynamic inter-lineage immune interactions. The events we documented within our model closely parallel mechanisms associated with checkpoint inhibitor-induced intestinal inflammation and inflammatory bowel disease, which were identified using primary patient samples (40, 46, 56). Aside from affording the possibility for real-time, dynamic observation of TRM-related processes and immune-epithelial interaction, our model provides the advantage of direct perturbation and hypothesis testing. Whereas multi-omic analyses of primary patient samples provide rich catalogs of differences between baseline and diseased states, the exact roles of differentially regulated parameters are difficult to ascertain without manipulating them in a relevant context. After demonstrating that IIOs capture known mechanisms of drug-induced inflammation and recapitulate well-described mitigation strategies, we use them to identify novel approaches for the management of TCB-mediated toxicities. Specifically, we find that blocking TRM motility through the Rho pathway helps dampen inflammation. Intriguingly, small molecules that target this pathway are being developed as fibrosis inhibitors for inflammatory bowel disease (61). Our data suggest that an additional benefit of inhibiting Rho signaling could be the quenching of T-cell-driven inflammation, by reducing the motility, contractility and ability of patrolling IELs to respond to TCR stimulation and engage with epithelial cells. Bearing in mind that our simple model recapitulates both the phenotypic outcomes and the multi-compartment cellular interactions that mediate them, IIOs can be instrumental in investigating tissue-resident immune responses in contexts far beyond drug-induced inflammation, including tumorigenesis, infectious and autoimmune diseases.

---

**Extended Data Figure 1.**
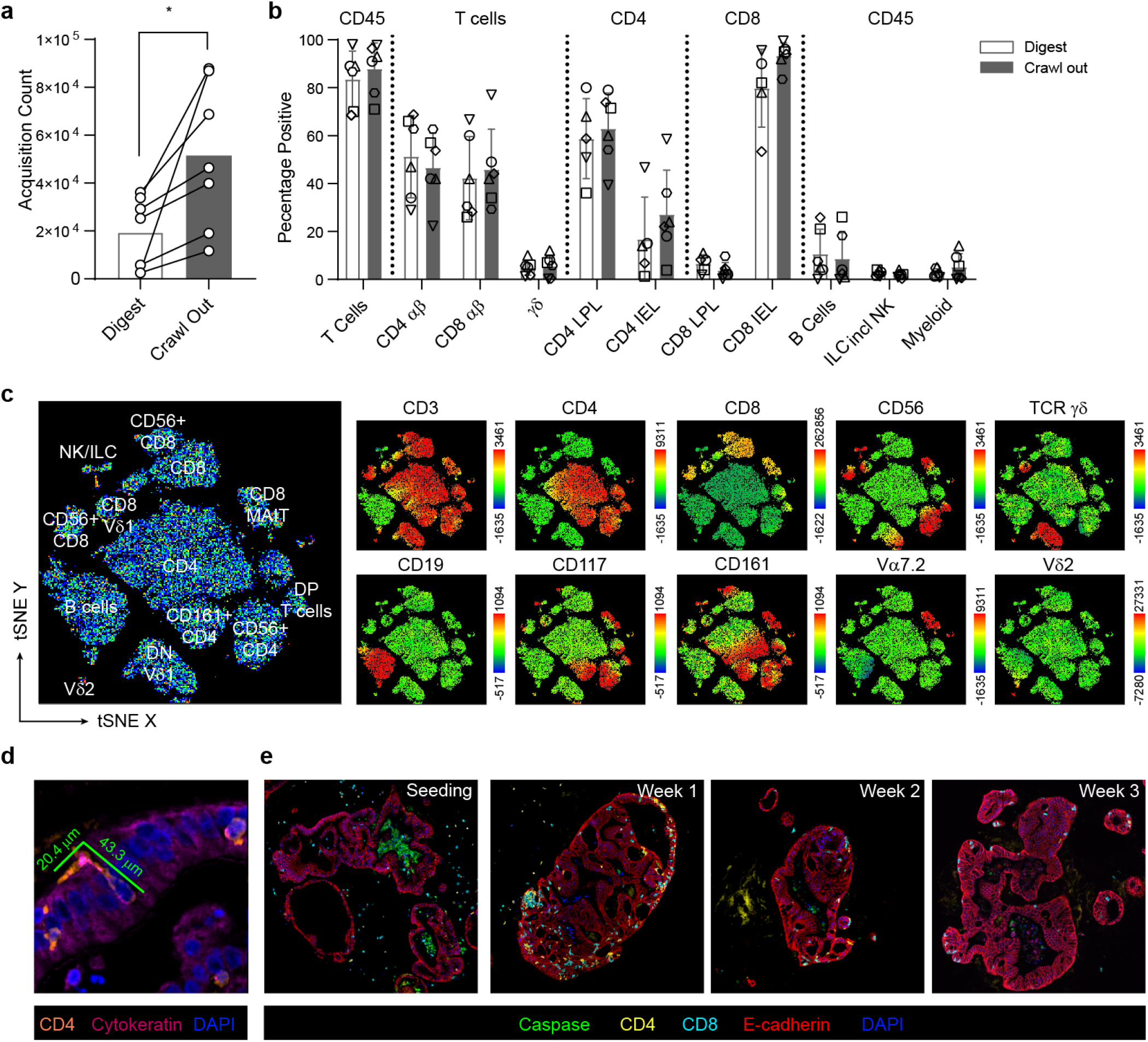
Intestinal TRM isolation and comparison to circulating T cells. a, Comparison of viable CD45 count in matched donors subjected to either digestion- or crawl out-based isolation. *P < 0.05, paired T-test. b, Comparison of immune cell proportions subjected to either digestion- or crawl out-based isolation. Values represent percentages of parent population listed above each bar. LPL lamina propria lymphocyte, IEL intraepithelial lymphocyte, DP double positive, DN double negative. c, tSNE analysis of a typical intestinal lymphocyte isolation incorporating standard lineage defining immune cell markers. Leftmost plot represents annotated cell populations with smaller plots displaying heatmaps for each of the 10 key surface markers assessed. d, Elongated flossing T cell inserting itself between basal-lateral epithelial cell junctions. Diameter of the T cell from head to tail is listed. e, Fluorescent IHC staining of the IIO culture over time.

**Extended Data Figure 2.**
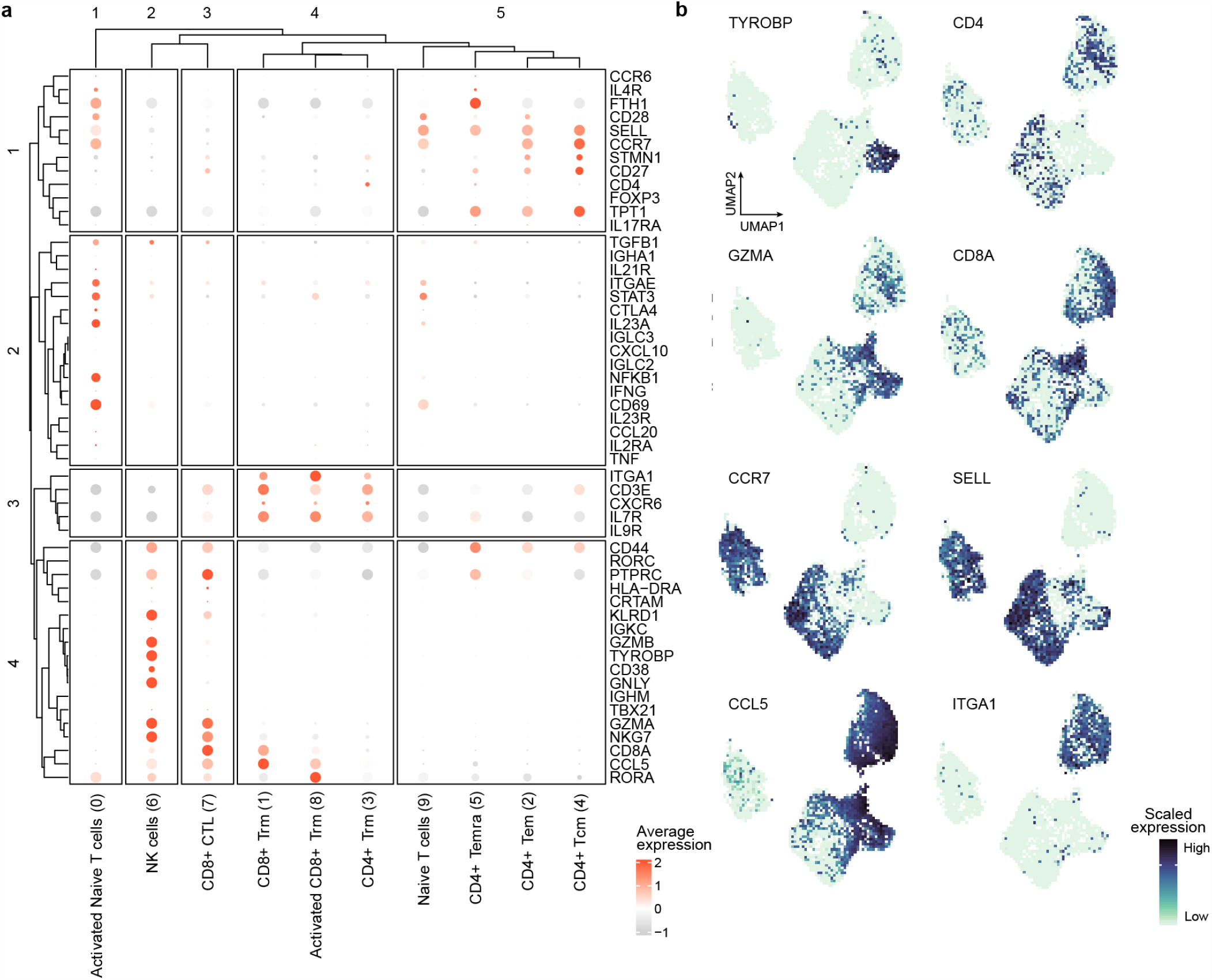
Immune cell cluster annotation. a, Dotplots showing the z-scored average expression of curated T cell subset marker genes for each cluster introduced in Fig. 2b. The size of the dot encodes the percentage of positive cells within a cluster for the respective gene. Subsets were classified into CD4+ and CD8+ populations using the aggregated expression values for CD4 and CD8A. Activated populations were identified using the expression of RORA (62). Tissue-residency of memory cells was determined based on the expression of ITGA1 and ITGAE, which are expressed by most TRMs (63). Final annotations were then designated based on the combinatorial expression of multiple markers. Source data detailing averaged expression values for T cell subset marker genes are provided in the source data file. b, Feature plot showing the scaled expression of representative marker genes.

**Extended Data Figure 3.**
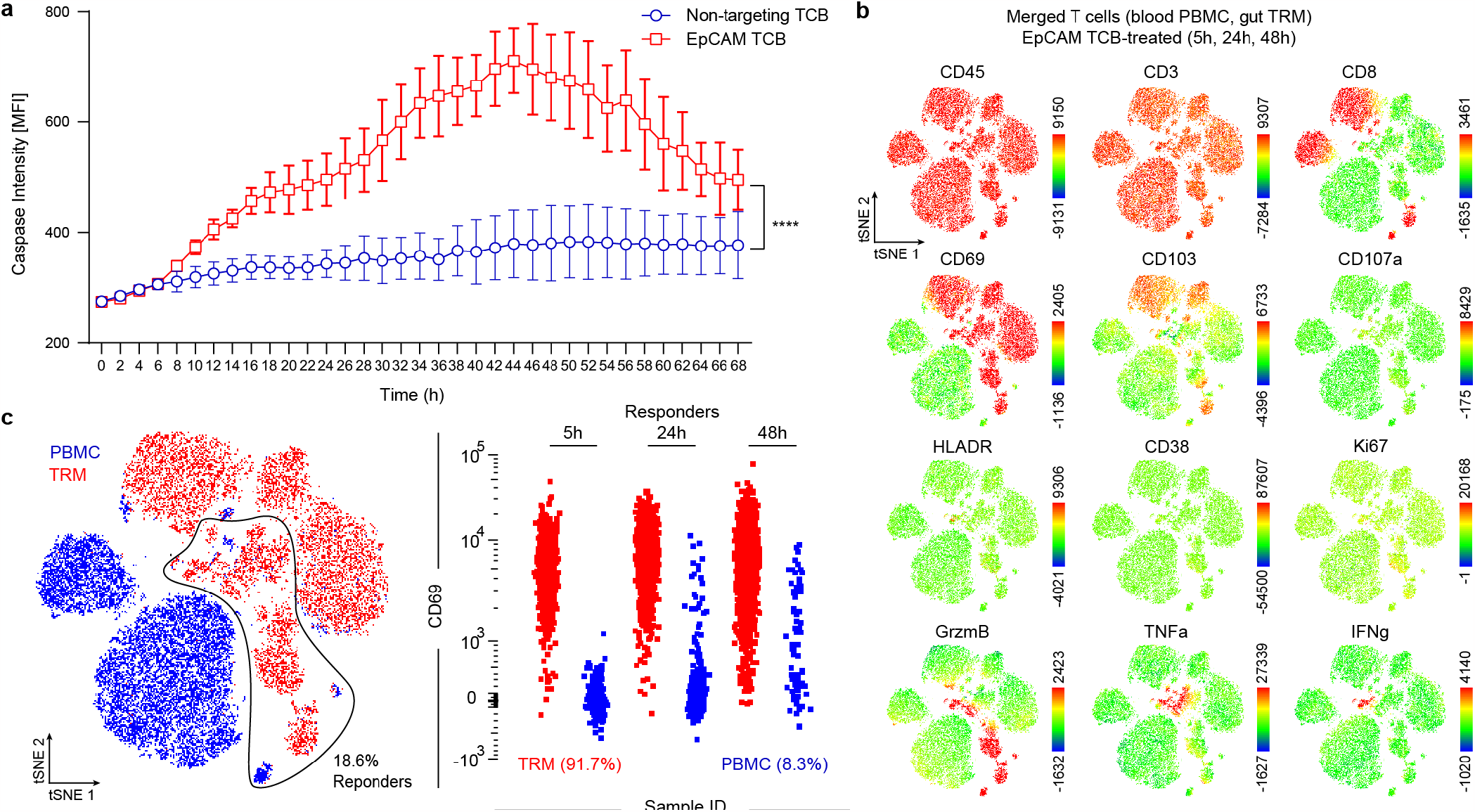
TRMs exhibit rapid and aggressive targeting of healthy epithelium following EpCAM-TCB treatment. a, bihourly quantification of caspase 3/7 signal in IIO cocultures for 68h treated with either 5 ng/ml EpCAM TCB or a non-targeting control molecule, n = 3, mean ± SD. ****P < 0.0001, T-test of the two AUCs. b, tSNE analysis of all T cells at all timepoints, derived from IIO cocultures with TRMs or PBMCs treated with 5 ng/ml EpCAM TCB. Plots display heatmaps for each of the 12 surface and intracellular markers assessed by flow cytometry. c, gating strategy for identifying responder cells (those that expressed TNFa, IFNy or GzmB), and the ungating of the concatenated flow cytometry files to reveal the original source of responder T cells.

**Extended Data Figure 4.**
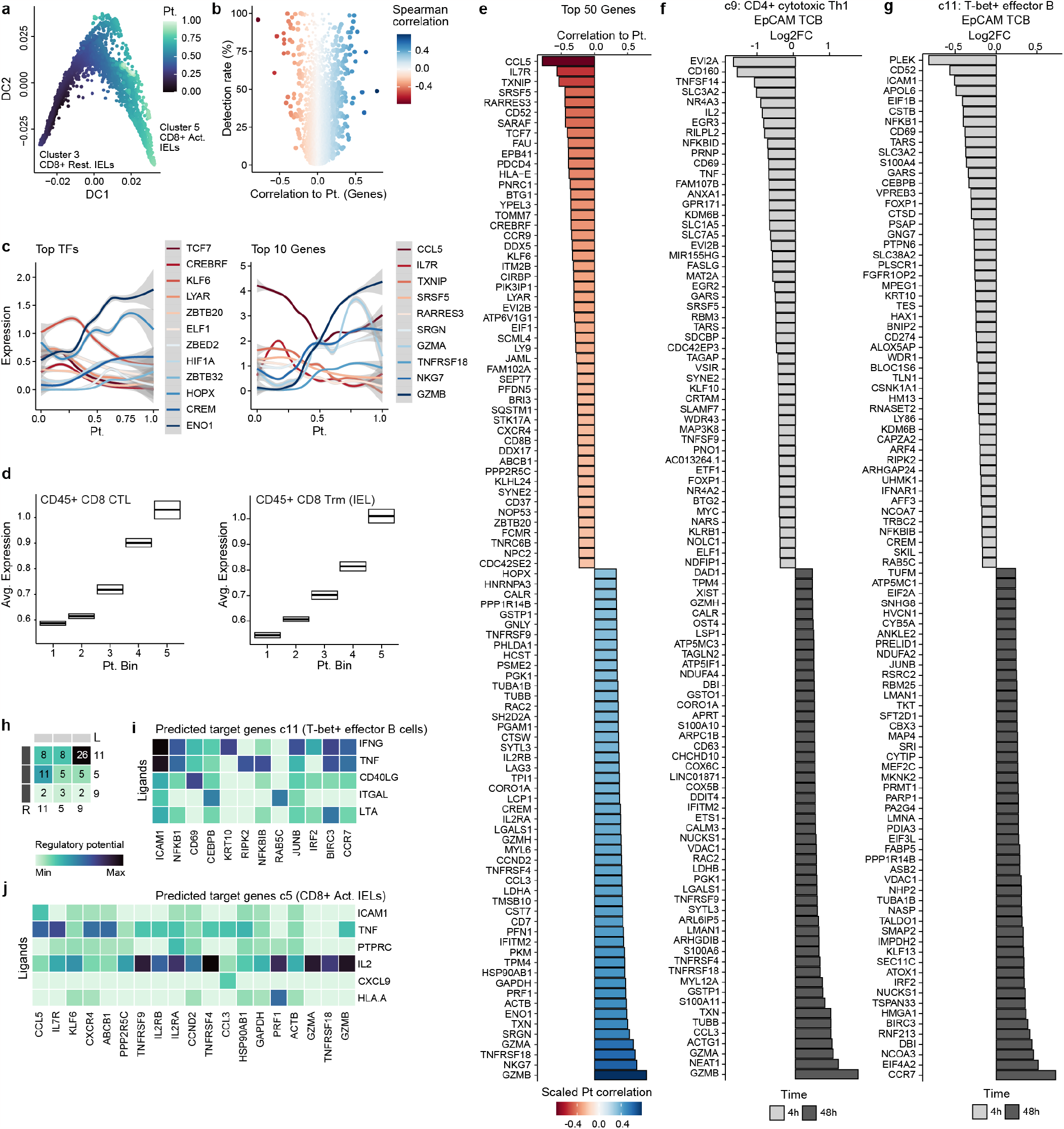
Extended analysis of cell states and interactions in IIOs after treatment. a, Plot represents IIO CD8+ T cells on the first two diffusion components colored by reconstructed diffusion pseudotime. b, Volcano plot showing variable genes along reconstructed CD8+ T cell trajectory. x-axis represents spearman correlation value between gene expression and pseudotime value, y-axis represents detection rates of genes in the CD8+ T cell populations. c, Expression patterns of most variable TFs (left) and effector genes (right) along the reconstructed CD8+ T cell trajectory. d, Boxplot summarizing the expression pattern of CD8+ CTL (left) and CD8+ Trm IEL (right) gene signatures described in ref 40 along the reconstructed activation trajectory of IIO CD8+ T cells. x-axis represents 5 equidistant bins of the reconstructed pseudotime. e-g, Barplot showing differentially expressed genes in the CD8+ T cell trajectory (e), in the CD4+ cytotoxic Th1 population (c9) (f) and in the T-bet+ effector B cells (c11) (g) at 4h and 48h after EpCAM TCB treatment. h, Heatmap summarizes directed ligand-receptor pairing interactions of T-bet+ effector B cells (c11) and T cell populations (clusters 5 and 9) in the IIO model. i-j, Predicted regulatory potential of CD4+ cytotoxic Th1 cells (c9) ligands towards signature genes of T-bet+ effector B cells (c11) (i) and CD8+ activated IELs (c5) (j).

**Extended Data Figure 5.**
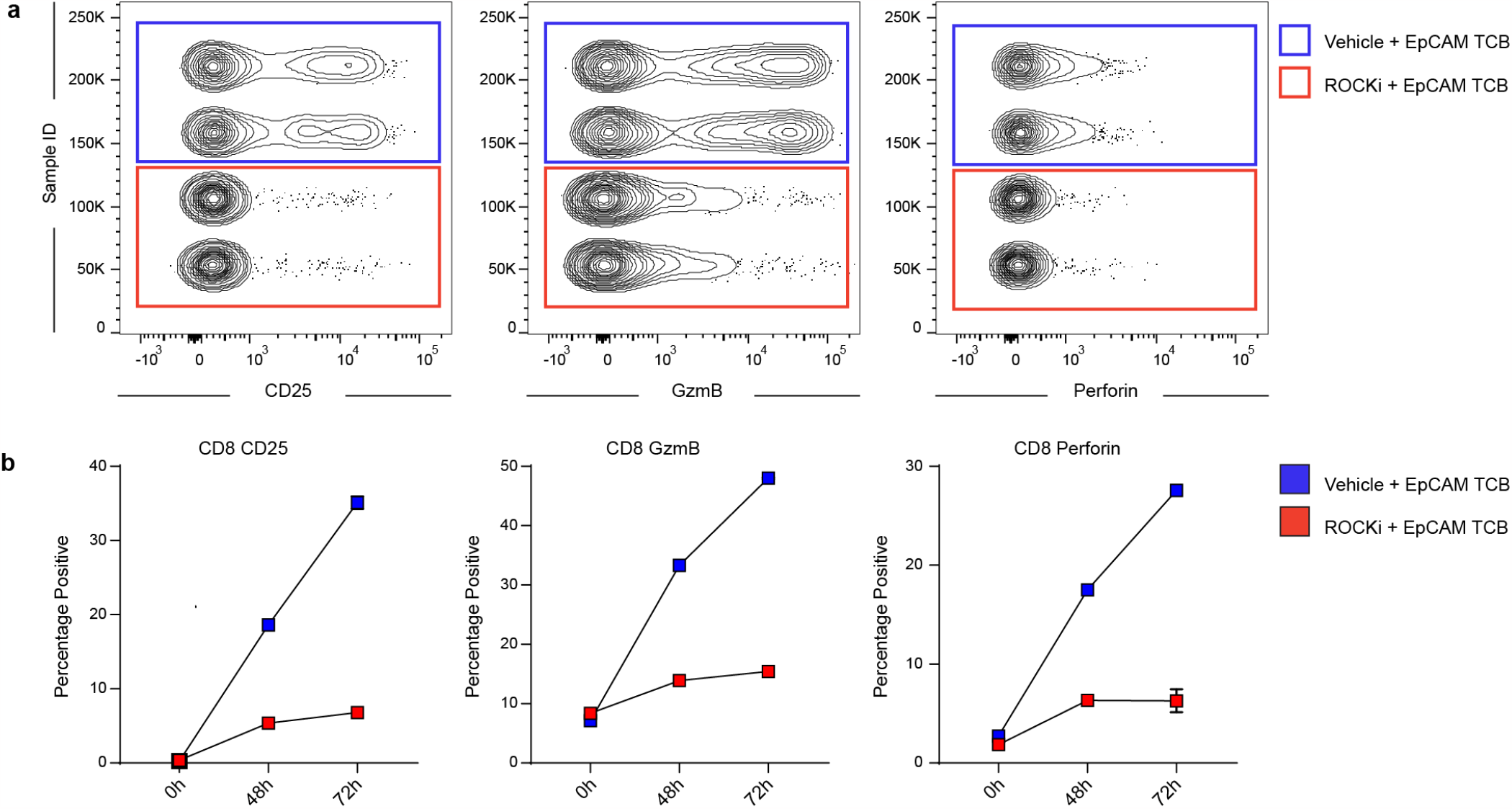
Inhibition of ROCK1/2 pathway quells TRM-driven intestinal inflammation. a, flow cytometry plots comparing expression of CD25, GzmB and perforin in CD8 T cells isolated from IIO cocultures 72h after treatment with 5 ng/ml EpCAM-targeting T-cell bispecific antibodies with or without 1 mM Y27362 (ROCKi) b, Line graphs quantifying expression of CD25, GzmB and perforin in CD8 T cells isolated from IIO cocultures at baseline, at 48h and at 72h after treatment with 5 ng/ml EpCAM-targeting T-cell bispecific antibodies with or without 1 mM Y27362 (ROCKi).

## Methods

### Isolation of tissue-resident memory cells and donor– matched crypts for organoid generation

Human intestinal tissue samples and annotated data were obtained and experimental procedures performed within the framework of the non-profit foundation HTCR (Munich, Germany) including informed patient’s consent. Tissue was first processed by removing the underlying muscularis, serosa and fat from the basal side of the tissue by pinching with forceps and trimming away with scissors. The remaining mucosal tissue was washed with PBS (supplemented with penicillin [1500U/ml], streptomycin [1500 pg/ml], gentamicin [500 pg/ml]) multiple times before using a scalpel to remove excess mucus from the luminal side and blood vessels from the basal side. Trimmed, cleaned tissue was then cut into square explants approximately 5 mm by 5 mm with a scalpel. Two methodologies were used to isolate intestinal tissue-resident memory cells. For the scaffold-based egression protocol (adapted from (29)) each explant was loaded onto a 9x9x1.5 mm tantalum-coated carbon based scaffold (Ultramet), and cultured in 24-well plates containing 1 mL of cytokine-free, cell culture media (RPMI 1640 10% FCS, penicillin [1500U/ml], streptomycin [1500 pg/ml], gentamicin [500 pg/ml] and amphotericin [12.5 pg/ml]). 24 hours later, scaffolds and tissue were removed, and egressed cells were harvested from the bottom of the culture well via pipetting. For the enzymatic digestion protocol, explants were digested using the Human Tumour Dissociation Kit with the gentleMACS Octo Dissociator program 37C_h_TDK_1 (Miltenyi Biotec), as per the manufacturer’s instructions, reducing Enzyme R content by 80% to minimize surface antigen cleavage. Cells were then counted, phenotyped by flow cytometry and frozen prior to use in downstream applications. To isolate donor-matched crypts for organoid generation, a lamelle was scraped over a 3 cm2 piece of trimmed tissue to remove intestinal protrusions/villi. Tissue was then incubated in ice cold PBS + 10 mM EDTA with vigorous shaking for 30 minutes to break down epithelial cell junctions. The tissue was retrieved and a lamelle was used to scrape off crypts into DMEM-F12 1% BSA collection buffer. Crypts were centrifuged, resuspended in Matrigel Matrix GFR (Corning) and cultured in IntestiCult™ Organoid Growth Medium with 10 uM Y-27362 (STEMCELL Technologies).

### Preparation and culture of intestinal immune-organoids

In-passage organoids, approximately 2 weeks to 1 month after initial isolation, were cultured for 4 days post-split in IntestiCult™ Organoid Growth Medium and then switched IntestiCult™ Organoid Differentiation Medium (STEMCell Technologies) for 72 hours to promote epithelial stem cell differentiation. On the day of coculture setup, media was aspirated from the well, organoids were washed with PBS and treated with ice cold Cell Recovery Solution (Corning) for 40 minutes. The solution was collected and centrifuged to harvest the liberated organoids. Donor-matched TRM or PBMCs were thawed, and combined with the liberated organoids and resuspended in either Matrigel Matrix GFR (Corning), or if the cocultures were to be formalin-fixed, a 50:50 mixture of Matrigel Matrix GF and 4 mg/ml Cultrex Rat Collagen I (R&D Systems). For time-lapse live imaging experiments, immune cells were labelled with CellTrace Far Red or CFSE (ThermoFisher) prior to mixing with organoids. Organoids were used at a concentration double to their standard passaging density, whereby a 20 ul dome was harvested and resuspended in 10 ul of matrix, while immune cells were used at a density of 15,000 cell per mm3 of resuspension volume. A 50:50 ratio of RPMI 1640 10% FCS and IntestiCult™ Organoid Differentiation Medium was used for culture. For assessment of TCB-based cytotoxicity, an EpCAM-CD3 bispecific antibody, or its associated non-targeting control (which contains a CD3 binder on one arm and a non-specific DP47 arm), was supplemented into the culture medium upon coculture setup, typically at a concentration of 5 ng/ml. When used, blocking antibodies to TNFa (MAb-1, Biolegend) were added at a concentration of 1 ug/mL. To investigate the role of ROCK signaling on TRM activation, 10 uM Y-27362 was added daily for the duration of the coculture. For the month-long cocultures, the media was supplemented with IL-2 [10 lU/mL] (Roche) and IL-15 [2 ng/ml] (BioLegend), media change 3 times per week and culture splitting 1 time per week. Cultures were supplemented with 10 uM Y-27362 (STEMCELL Technologies) after splitting.

### Flow cytometry analysis of immune-organoids

Triplicate wells of immuno-organoid cocultures from each condition were harvested 5h, 24h and 48h post-treatment. 4h prior to culture harvest at each timepoint, wells were treated with Protein Transport Inhibitor Cocktail (Thermo Fisher) to facilitate intracellular accumulation of cytokine protein. Co-cultures were washed with PBS and then digested to single cells using Accutase solution (STEMCELL Technologies) at room temperature for approximately 30 minutes. Cell suspensions were passed through a 70 um strainer and stained for surface proteins (Extended Data Table 3). Cells were then fixed and permeabilized using the Foxp3 Transcription Factor Staining Buffer Set (Thermo Fisher), and subsequently stained for intracellular and intranuclear proteins. Stained cell suspensions were acquired on a BD Fortessa X-20 and analysed using FlowJo v10. To facilitate visual representation across one plot, samples from different time points and treatments were concatenated and separated along the y-axis.

### Caspase-3/7 based epithelial cell cytotoxicity assay

IIO cultures were prepared in 4 ul Matrigel Matrix GF per well of a PhenoPlate™ 96-well microplate (Phenoplate), with cell ratios and media as described above. Apoptosis was assessed using the CellEvent Caspase-3/7 Detection Reagent (Invitrogen) over TCB-treatment duration or at specific intervals. CellEvent Caspase-3/7 Detection Reagent was to the culture media at 1:1000. Samples were imaged in confocal mode at 5x magnification (Air objective) with the Operetta CLS (Perkin Elmer) covering approximately a 450 pm Z-stack, starting at -150 pm. Distance between Z-stacks was set to the minimum of 27 pm for the 5x objective (Autofocus: Two Peak; Binning: 2). Channels selected were brightfield and the predefined AlexaFluor 488. Per well, 5 fields were acquired, covering nearly the entire surface of the 96-well PhenoPlate plate. CO2 was set to 5%, temperature to 37°C. Caspase 3/7 fluorescence signal intensity was quantified using the Opera Harmony software (PerkinElmer). Briefly, segmentation of organoids was done by using “Find Texture Regions” based on the bright-field signal only, followed by “Select Region” and “Find Image Region” to segment single organoids as objects. Next, “Calculate Morphology Parameters” was performed to select objects >10000 pm 2 with “Select Population”. Next, the Caspase 3/7 fluorescence signal per individual organoid was determined using “Calculate Intensity Properties” of the AF 488 channel within these objects.

### Time-lapse imaging of IIOs

The time-lapse live imaging was performed with a Leica SP8 confocal microscope using a 20x dry objective (HC PL APO CS2 20x/0.75 DRY) and 0.75 zoom. A 119.93pm thick Z-stack (step size: 3.98pm) was imaged every 10 minutes. During imaging the samples were in an incubation chamber (The Box, Life Imaging Services) at 37°C and 5% CO2. After acquisition maximum intensity projections were generated with the Leica Las X software and later exported as AVI files using ImageJ.

### FFPE-embedding of cocultures

To FFPE-embed the co-cultures, the samples were seeded in a 50% (v/v) Matrigel-Collagen I matrix. Wells were washed once with 1X DPBS before fixation with 4% paraformaldehyde (PFA) in the 24-well Clear TC-treated plate. After 30 min of fixation at RT, the wells were washed three more times before complete aspiration of the 1X DPBS. 400 pL preliquefied HistoGel (Thermo Scientific) was dispensed into the 24-well Clear TC-treated plates. After polymerization of the HistoGel (10min in 4°C fridge), the organoid-HistoGel ‘platelet’ was carefully lifted out of the 24-well Clear TC-treated plate using a thin metallic spatula. The samples were then distributed into biopsy cassettes and dehydrated overnight using a Vacuum filter processor (Sakura, TissueTek VIP5). On the next day, the samples were embedded in liquid paraffin.

### Microtome sectioning

FFPE blocks were - in general - sectioned at a thickness of 5 pm and transferred on Super-frost Plus Adhesion microscope slides (Epredia). Where indicated, thickness differs. Slides were incubated in a slide oven overnight at 37°C.

### Staining - FFPE-based multiplex immunofluorescence (mIF)

mIF staining of FFPE slides was performed using Ventana Discovery Ultra automated tissue stainer (Roche Tissue Diagnostics, Tucson AZ USA). Slides were baked first at 60°C for 8 min and subsequently further heated up to 69°C for 8 min for subsequent deparaffinization. This cycle was repeated twice. Heat-induced antigen retrieval was performed with Tris-EDTA buffer pH 7.8 (Ventana) at 92°C for 32 min. After each blocking step with Discovery Inhibitor (Ventana) for 16 min, the Discovery Inhibitor was neutralized. Primary antibodies were diluted in 1X Plus Automation Amplification Diluent (Akoya Biosciences). Primaries were detected using according anti-species secondary antibodies conjugated to HRP (OmniMap Ventana) (Extended Data Table 3). Subsequently, the relevant Opal dye (Akoya Biosciences) was applied. After every application of a primary, respective secondary antibody and opal dye, an antibody neutralization and HRP-denaturation step was applied to remove residual antibodies and HRP, before starting the staining cycle again with the Discovery Inhibitor blocking step. Lastly, samples were counterstained with 4’,6-Diamidino-2-phenylindol (DAPI, Roche).

### FFPE-based mIFe

mIF stainings using the Opal dyes from Akoya were digitized with multispectral imaging by the Vectra® Polaris™ (PerkinElmer) using the MOTiF™ technology at 20x magnification for all 7 colors (Opal 480, Opal 520, Opal 570, Opal 620, Opal 690, Opal 780 and DAPI). Slides were scanned in a batch manner to ensure same imaging settings and cross-comparability for later image analysis with HALO AI. Next, unmixing of the channels and tiling of the images was performed with PhenoChart (v1.0.12) and inForm (v2.4). Tiles were fused in HALO (Indica labs, v3.2.1851.328). High-resolution mIF were obtained using a STELLARIS 8 microscope (Leica) with a 40X1.1 NA water-immersion objective (HC FLUOTAR L VISIR 25x/0.95 WATER) and 1.0 zoom. The white light laser (WLL: 440 nm-790 nm) allowed to image all opal dyes mentioned above, channels were acquired sequentially to reduce crosstalk. Images were obtained in bidirectional mode with 2048 x 2048 pixels (pixel size: 273.8nm x 273.83 nm) at 600 Hz. Where indicated, images were acquired with z-stacks between 10-15 pm (z-steps: 1 pm) and 3D-reconstructed and displayed in ‘Maximum’-mode using the Leica Application Suite X software (Leica).

### Image Analysis - FFPE-based mIF

Image analysis of the mIF images was performed with HALO AI (Indica Labs, v3.2.1851.328). Briefly, single organoids were automatically detected using a Deep Learning algorithm trained to distinguish matrix and organoids (iterations: 5000; cross entropy: 0.32; DenseNet AI V2 Plugin). After quick validation, organoids were annotated as individual ROIs, objects. Only objects >7500 pm2 were considered positive. The HighPlex FL v4.1.3 module was used to perform nuclear segmentation based on DAPI+ cells. For quantification, DAPI+ nuclei and markers for each distinct cell type of interest were merged. Thereby, secondary-only negative controls on the tissue of origin served as a negative signal threshold in order to prevent biased adjustments. The analysis module was deployed on ROIs per single object (organoid).

### Single-cell dissociation of IIO

IIO were dissociated as described previously by (64). In short, organoids were dislodged and mechanically broken up and transferred to 1% BSA-coated tubes. The organoid fragments were centrifuged at 400g, 4min, RT. The supernatant was removed and the enzymes of the neural tissue dissociation kit (Miltenyi Biotec) were mixed in HBSS-1%BSA buffer. The organoid fragments were dissociated to single cells for a total of 30 min with through pipetting every 7 minutes. Next, cells were filtered through a 40uM filter and single cells were centrifuged at 450g, 4min, RT and subsequently resuspended in DPBS 1% BSA. The single cell libraries were prepared with the 10x Genomics platform using the Chromium Next GEM Single Cell 3’ Kit v3.1.

### Single-cell RNA-seq data preprocessing

CellRanger (v6.0.2, 10x Genomics) was used to extract unique molecular identifiers, cell barcodes, and genomic reads from the sequencing results of 10x Chromium experiments. Then, count matrices, including both protein coding and non-coding transcripts, were constructed aligning against the annotated human reference genome (GRCh38, v3.0.0, 10x Genomics). In order to remove potentially damaged or unhealthy cells and improve data quality, the following filtering steps were performed in addition to the built-in CellRanger filtering pipeline. Cells associated with over 50,000 transcripts, usually less than 1% of the total number of samples, were removed. Cells associated with a low number of unique transcripts, less than 500 unique transcripts detected (1% of the total number of samples), were removed. Cells with over 20% of mitochondrial transcripts were removed. Transcripts mapping to ribosomal protein coding genes as well as mitochondrial genes were removed together with transcripts detected in less than 10 samples.

### Normalization with SCTransform

For normalization and variance stabilization of each scRNA-seq experiment’s molecular count data, we employed the modeling framework of SCTransform in Seurat v3 (65). In brief, a model of technical noise in scRNA-seq data is computed using ‘generalized gamma poisson regression’ (66). The residuals for this model are normalized values that indicate divergence from the expected number of observed UMIs for a gene in a cell given the gene’s average expression in the population and cellular sequencing depth. Additionally, a curated list of cell cycle associated genes, available within Seurat, was used to estimate the contribution of cell cycle and remove this source of biological variation from each dataset in order to increase the signal deriving from more interesting processes. The residuals for the top 2,000 variable genes were used directly as input to computing the top 100 Principal Components (PCs) by PCA dimensionality reduction through the RunPCA() function in Seurat. Corrected UMI, which are converted from Pearson residuals and represent expected counts if all cells were sequenced at the same depth, were log-transformed and used for visualization and differential expression (DE) analysis. Primary intestinal biopsy samples as well as primary multiorgan biopsy samples were processed as described above. However, they did not undergo any cell filtering as quality control steps had already been performed in the respective published studies.

### Doublet removal with DoubletFinder

For each scRNAseq experiment DoubletFinder (67) was used to predict doublets in the sequencing data. In brief, this tool generates artificial doublets from existing scRNA-seq data by merging randomly selected cells which are then pre-processed together with real data and jointly embedded on a PCA space that serves as basis to find each cell’s proportion of artificial k nearest neighbors (pANN). For this step we restricted the dimension space to the top 50 PCs. Finally, pANN values are rank ordered according to the expected number of doublets and optimal cutoff is selected through ROC analysis across pN-pK parameter sweeps for each scRNA-seq dataset; pN describes the proportion of generated artificial doublets while pK defines the PC neighborhood size. In order to achieve maximal doublet prediction accuracy, mean-variance normalized bimodality coefficient (BCmvn) was leveraged. This provides a ground-truth-agnostic metric that coincides with pK values that maximize AUC in the data. To overcome DoubletFinder’s limited sensitivity to homotypic doublets, we consider doublet number estimates based on Poisson statistics with homotypic doublet proportion adjustment assuming 1/50,000 doublet formation rate the 10x Chromium droplet microfluidic cell loading.

### Ambient mRNA signal removal

After doublet prediction and removal, we analyzed each scRNA-seq dataset in order to estimate the extent of ambient mRNA contamination in every single cell and correct it. We used the R package Cellular Latent Dirichlet Allocation (CELDA) (68) which contains DecontX, a method based on Bayesian statistical framework to computationally estimate and remove RNA contamination in individual cells without empty droplet information. We applied the DecontX() function in CELDA to the raw count matrices with default parameters. Subsequently, we removed all cells with contamination values above 0.5 and we used the decontaminated count matrices resulting from DecontX() for downstream analysis.

### Data integration

Individual datasets, after preprocessing, doublet removal and ambient mRNA regression, were aggregated according to specific criterias (e.g. tissue of origin, profiling time, culture condition) and went through a joint normalization step with SCTransform in order to mitigate technical confounding factors, which also served as means for selection of a set of meaningful 2,000 most variable global genes prior to data integration. Integration of different conditions (culture model, treatment and timepoints) was performed using the log-normalized corrected UMI count data in two steps. First, the residuals for the top 2,000 global variable genes were used as input to computing the top 100 Principal Components (PCs) through the RunPCA() function in Seurat. The leading 30 PCs and 50 nearest neighbors were then used to define the shared neighborhood graph with the FindNeighbors() function in Seurat. Subsequently, datasets were clustered according to the shared neighborhood graph using the Louvain algorithm (69) through the Seurat function FindClusters() with resolution 0.8. Finally, we used these high-resolution clusters to define a restricted, noise reduced and cell state specific set of genes through differential expression analysis (see Methods section below). In the second step of the integration process, we compiled a list consisting of the ensemble of the top 30 DEGs for each cluster and used it to focus and repeat PCA dimensionality reduction. The first 30 PC vectors of the new PCA space served as basis to obtain a two-dimensional (2D) representation of the data through Uniform Manifold Approximation and Projection (UMAP) (70) implemented in RunUMAP() with 50 nearest neighbors. We then computed a shared neighborhood graph on the UMAP lower-dimensional space and computed the final integrated clusters with resolution parameter 0.2.

### Differential expression analysis

Gene differential expression (DE) analysis between distinct cell populations in scRNA-seq data was assessed by performing Wilcoxon rank sum tests and auROC analysis as implemented by Presto (71) package in R. Log-transformed corrected UMIs were used as input for the DE statistical tests, and genes were called differentially expressed if associated adjusted p-value (Bonferroni method) was lower than 0.05, AUC value was above 0.6 and log fold change was greater than 0.15. In addition, we also set thresholds on detection rates of DE genes. In particular, a given gene was assigned as over-expressed in the analyzed group if it was detected in at least 30% of the samples of that group, while the detection rate in the background samples was at most 70% of the detection rate of the analyzed group. CD8+ T cell activation trajectory reconstruction. To reconstruct the continuum of the CD8+ T cell activation trajectory in IIO models challenged with bispecific antibodies we took advantage of diffusion pseudotime (DPT) as implemented by the destiny package In R. In brief, DPT uses random-walk-based distance, computed on the leading eigen-vectors of a transition matrix, to order scRNA-seq data according to differentiation stages (49, 72). Concretely, we used the DiffusionMap() function in destiny package (73) on the space identified by the leading 30 PC vectors of the integrated PCA embedding of the CD8+ clusters. Pseudotime values were then computed with the DTP() function in destiny on the diffusion map object using default parameters. Similarly, the global pseudotime, following TNF perturbation simulation, was based on a random walk approach on the cell state transition matrix.

### Intercellular communication analysis

To investigate ligand-receptor (LR) mediated cell-cell communications during immune cell activation in our IIO models, we focused on the signals exchanged between Th1 cells, activated t-bet B cells and CD8+ CTLs. For this analysis we extracted genes labeled as either ligands or receptors from curated databases A.and required that genes were differentially expressed between the three populations under investigation, which facilitated retrieval of directional information about the signal exchange. To gain insights into functional cell-cell communication, we used the NicheNet pipeline which considers the influence of sender-cell ligands on receiver-cell gene expression (58). NicheNet’s analysis pipeline provided us with a ranking of predicted ligands that most likely affect gene expression in activated t-bet B cells and CD8+ CTLs highlight-ing the role of critical Thl-secreted factors in driving immune cell phenotypes within IlOs.

### Functional enrichment analysis

To understand mechanisms underlying phenotypes in our data, differentially expressed genes were analyzed for gene ontology biological process (GOBP) enrichment using one-sided hypergeometric testing. P-values were adjusted for multiple testing hypotheses by the Bonferroni method and only enrichment results below a 5% significance level threshold were considered. For this analysis, we only considered biological processes consisting of sets with more than 10 but less than 300 mapped genes.

### In silico perturbation analysis

To simulate dynamic shifts in cell identity resulting from ligand signaling cascade activation, we employed Nichenet’s prior model (58). The first step involved generating simulated values by applying the gene regulatory network (GRN) as a function and propagating the relative changes in gene expression after k-nearest neighbor imputation of the gene expression data. This iterative (3 times) signal propagation enabled us to calculate the broad, downstream effects of ligand perturbation, thereby estimating the global transcriptional shift. The estimation of cell-identity transition probability was accomplished by comparing this gene expression shift to that of local neighbors, utilizing a likelihood-based dynamical model. By doing so, we could establish a measure of how cell identities transition in response to ligand perturbation. Finally, the transition probabilities were transformed into a weighted local average vector map, encoding the simulated directionality of cell-state transition for each cell. This workflow results from an adaptation and integration of CellOracle (74) and scVelo (75).

